# Disentangling blade and vasculature shape in grapevine leaves

**DOI:** 10.64898/2026.01.27.701982

**Authors:** Wahiba Yahiaoui, Safa Smail, Yemeen Ayub, Qiao Lu, Peter Cousins, Luis Diaz-Garcia, Margaret Frank, Lauren Headland, Ciera Martinez, Zoë Migicovsky, Aashish Ranjan, Neelima Sinha, Joel F. Swift, Efrain Torres-Lomas, Elizabeth Munch, Ziane Laiadi, Daniel H. Chitwood

## Abstract

The leaf blade and vasculature develop together within a shared morphological space. Despite shared molecular patterning pathways, it is unknown if developmental and evolutionary variation affect these tissues separately or together in a coordinated way. Grapevine leaves have a morphometric history and abundant data measuring the shape of the blade and vasculature together. Using a combination of topological data analysis and deep learning, we perform reciprocal semantic segmentation of leaf blade and vasculature. Each tissue contains sufficient information to predict the other. We hypothesize that this is due to a one-to-one relationship between blade and vein. Using thin plate splines to swap and warp different combinations of blade and vein shapes, we show that a set of leaves with a many-to-one relationship of blade and vein are distinguishable from true leaves. We also swap blade and vein across the developmental series and between species and show that only reversing the developmental series disrupts the relationship between blade and vasculature. We end by discussing the evolutionary and developmental implications that there is a unique, one-to-one mapping between blade and vein that allows each to be predicted from the other.

**Author summary:** Leaves are made of two closely connected parts: the flat blade that captures light and the network of veins that transports water, nutrients, and developmental signals. Although these tissues grow together and share common molecular patterning pathways, it has remained unclear whether a particular blade shape is uniquely linked to a specific vein pattern. In this study, we use grapevine leaves as a model system and combine mathematical shape analysis with deep learning to examine this relationship. We show that the shape of the blade alone can accurately predict the vein network, and that the vein network can likewise predict the blade. This finding suggests a near one-to-one relationship between these two tissues. To test this idea, we created artificial leaves in which blade and vein shapes were deliberately mismatched. Although these synthetic leaves appeared realistic at a global level, a neural network was able to distinguish them from real leaves based on subtle differences. We further show that this tight coupling is maintained by the developmental sequence of leaf growth rather than by species identity, revealing a conserved constraint linking leaf form and internal structure.

## 1 Introduction

The blade and vascular system of the leaf exist in a shared, constrained space. Physically, these two tissues are embedded in a single leaf with a single morphology. Especially for the case of primary and secondary veins, the direction of the vasculature often points towards or terminates at the margin (or towards marginal veins, when present) coinciding with serrations, lobes, leaflets, and other marginal features. The coincident features of blade and vasculature represent overlapping, shared developmental mechanisms. The adaxial-abaxial patterning common to the blade and vasculature arises in the incipient leaf and precedes the outgrowth of the primordium that it initiates (1). Auxin not only presages the incipient leaf and leaflet location (2–5), but also where the leaf blade will expand. Auxin flux also defines where the future procambium will differentiate (6). From early stages of leaf initiation, auxin-mediated patterning links blade outgrowth and vascular differentiation within the leaf primordium. That the leaf blade and vasculature are constrained to the same physical space and that the developmental pathways that pattern each are shared between them is established. However, it remains an open question if there is a unique mapping between the patterning of the blade and vasculature, if a specific patterning of the vasculature dictates a unique form of the blade, and vice versa.

Historically grapevines (*Vitis* and *Ampelopsis* species) are a unique system to study the geometry of the leaf blade and vasculature. Motivated by questions of provenance and appellation, early ampelographers mathematically studied the shape of leaves to differentiate species and varieties. All grapevine leaves contain five major veins, which standardizes geometric comparisons between leaves. Both the angles of the primary and secondary veins and the shape of the blade, notably the depth of the lobes, were used to distinguish varieties (7–14). Later, mathematical models that could average leaves while preserving their local features were developed (15). The geometric framework of ampelography paved the way for detailed morphometric studies using Procrustean landmark-based approaches that have extensively sampled genetic differences in leaf shape between *V. vinifera* varieties (16) and developmental and environmental differences in leaf shape among North American wild species (17–19). Recently the Procrustean landmark-based approach was extended to represent the outlines of the blade and vasculature at high resolution with methods to sample the underlying morphospace to create synthetic leaves that can be used in training deep learning models (20,21).

Quantitative representations of leaf shape have long been used to infer the genetic, developmental, and environmental processes that pattern leaves. In grapevine, morphometric descriptions of blade outlines and venation geometry capture biologically meaningful variation across species, development, and environment. In this framework, both the leaf blade and vasculature can be treated as coupled geometric representations of a single developmental outcome, allowing their relationship to be analyzed quantitatively. New image-based deep learning methods when combined with geometric morphometric approaches allow previously intractable questions to be answered. Using these high-resolution representations, a thin plate spline representation (22) can be used to warp any blade form to that of the vein, or vice versa, by using the shared, coincident margin points as anchors. Convolutional Neural Networks (CNNs) are a recent and powerful deep learning method for image data. While Procrustean morphometrics can generate a large number of synthetic samples necessary for training CNNs (21), there remains the problem that geometric morphometric representations are distilled shape information different from a simple shape mask (23,24). Recently we used a radial version of the Euler Characteristic Transform (ECT) that topologically featurizes the embedded graph of a closed contour that spatially corresponds with the original shape mask, permitting semantic segmentation tasks (25).

Here, we examine and unconfound the relationship between closed contour representations of the blade and vasculature in grapevine leaves. Training a semantic segmentation model on the radial ECT representation and shape mask of either the blade or vasculature, we show that a pixel representation of the other can be predicted. As either the blade or vasculature contains sufficient information to predict the other, we hypothesize that there is a one-to-one mapping between them. To test this hypothesis, we use thin plate splines to artificially warp many blade contours to a single vasculature and vice versa, creating datasets with “many-to-one” relationships between blade and vasculature. A two tower CNN that creates a common embedding space between true matches and the artificially created many-to-one leaves demonstrates a one-to-one mapping between blade and vasculature in real leaves. Finally, we examine leaf series across the first four leaf nodes of wild *Vitis* spp., swapping blade and vasculature between species or reversing the developmental series. A two tower CNN discriminates against artificially created series where the developmental progression has been reversed. Together, our results demonstrate the inextricable link between blade and vasculature in leaves, such that each contains sufficient information to predict the other, consistent with a near one-to-one relationship. Beyond differences in leaf shape among species, our results further indicate that a conserved developmental progression underlies the shapes of both blade and vasculature.

## 2 Materials and Methods

### 2.1 Synthetic data generation and Euler Characteristic Transform (ECT)

Previously published leaf coordinates from *Vitis* and *Ampelopsis* species (grapevines) representing domesticated varieties, rootstock, wild species, and dissected leaf type classes (21) were combined with new data collected from Algerian varieties (26) and were aligned via Procrustes analysis in a common Principal Component Analysis (PCA) morphospace using Python. Synthetic samples were generated in this PCA space using a Synthetic Minority Oversampling Technique (SMOTE)-like augmentation technique. This method interpolates between existing class members to inversely sample the morphological manifold, thereby creating novel, yet biologically plausible, leaf coordinates. These synthetic PCA scores were then projected back into the original high-dimensional coordinate space using the inverse PCA transformation to retrieve synthetic outlines. For each coordinate set, the ECT was computed for both the blade and vein components. The raw coordinates were first subjected to a standard set of geometric normalizations (centering, rotation, and scaling to a fixed radius=1) using the ECT module (https://munchlab.github.io/ect/). The radial ECT was calculated by conversion to polar coordinates and aligned to the original shape mask by using the affine transformation matrix derived from the normalization process.

### 2.2 Reciprocal U-Net Segmentation

In order to predict the vasculature using only blade information and vice versa, two separate U-Net models for reciprocal semantic segmentation on a synthetic leaf dataset were used. Each model utilizes two input channels (radial ECT and shape mask) and predicts a single-channel binary mask. The Blade-to-Vein model maps the blade ECT and blade mask (inputs) to the vein mask (target), while the Vein-to-Blade model maps the vein ECT and vein mask (inputs) to the blade mask (target). Training was conducted using the PyTorch framework (27), with a Combined Loss Function consisting of 50% Binary Cross-Entropy (BCE) and 50% Dice Loss, optimized with the Adam optimizer (learning rate: 0.0005) over 100 epochs. Training samples (approx. 80%) and validation samples (approx. 20%) were determined by a reproducible 42 seed split. For segmentation performance evaluation, the models were compared based on the Dice Coefficient calculated on the validation set after applying a 0.5 probability threshold to the sigmoid-activated outputs. Final trained models and checkpoint states were saved using PyTorch’s internal format (.pth), and model convergence was documented by saving the validation Dice history to a CSV file for comparative visualization, including both raw and Min-Max normalized scores.

### 2.3 Two tower CNN testing many-to-one relationships

To train the deep learning model to generalize across uncoupled morphological variation, two auxiliary synthetic datasets were created using Thin Plate Spline (TPS) interpolation. The TPS method relied on a set of 25 corresponding landmark points between shapes to define non-linear warping transformations. For any given source leaf and target, the target shape was first Procrustes-aligned to the source using their combined coordinates. The landmarks of the target were then iteratively warped to precisely align with the landmarks of source using an iterative TPS transformation, forcing landmark correspondence at the common landmark points between blade and veins. This resulting TPS function was then applied to all coordinate points of the target to generate the final warped shape. This procedure yielded two synthetic datasets: “warped blade” data, where the vein network of a leaf A was fixed and the blade outline of a different leaf B was morphed onto A’s vein landmarks; and “warped vein” data, where the blade outline of a leaf A was fixed and the vein network of a different leaf B was morphed onto A’s blade landmarks. These three datasets—matched, warped blade, and warped vein—were the basis for training and evaluating generalization to uncoupled variance between vein and blade outlines. A Two-Tower Convolutional Neural Network (CNN) was implemented for the cross-modal shape matching task, taking separate blade image and vein image inputs (composed of ECT and corresponding shape mask) derived from the normalized coordinates. The architecture employed two distinct CNN towers with identical internal structures, each designed to map its modality into a shared, low-dimensional latent embedding space of dimension 64. The model was trained end-to-end using the Contrastive Loss function, configured to minimize the distance between embeddings of matched pairs and maximize the distance between unmatched pairs. The training regime utilized a combination of the matched baseline data and the two synthetic warped datasets. Model performance was evaluated on the test set using standard retrieval metrics: mean Average Precision (mAP) and Recall@K for K= 1, 5, and 10, across both retrieval directions of blade-to-vein and vein-to-blade. The final model weights and the complete set of learned embeddings were saved for subsequent analysis. The effectiveness of the Contrastive Loss was quantified by analyzing the Euclidean distance between matched blade and vein embeddings and the global differences between groups of synthetic leaves visualized using Principal Component Analysis (PCA). The distribution of the resulting PC scores across the three data sources was compared using the Kruskal-Wallis test followed by Benjamini-Hochberg (BH) correction (alpha=0.05) to identify statistically significant changes in specific morphological components caused by the warping. Linear Discriminant Analysis (LDA) was applied to the PC scores to assess the linear separability of the data sources in the morphological space. The structure of the latent space was visually inspected using t-distributed Stochastic Neighbor Embedding (t-SNE).

### 2.4 Two tower CNN comparing inter- and intraspecies swaps

Leaves from the first four nodes of the tip of the shoot for ten different *Vitis* spp. (*Vitis acerifolia, V. aestivalis, V. amurensis, V. cinerea, V. coignetiae, V. labrusca, V. palmata, V. riparia, V. rupestris, V. vulpina*) were considered together as a stacked unit. For each blade and vein contour for each leaf in the series a shape mask and corresponding radial ECT were calculated. For intraspecies swaps, the order of blade or vein contours across the four nodes was reversed relative to the other within each species. For each swap, warping of blade-to-vein and vein-to-blade were performed. As the order of blade to vein is swapped across the series and each of two possible warpings is performed, we took each possible ordering of the series. Ultimately, we found a difference between orderings that we term “with the series” or “against the series”. “With the series” is defined as the unwarped component retaining the true node order; “against the series” is defined as the unwarped component ordered in the reverse direction of the true direction. A Two-Tower CNN was implemented for the cross-modal retrieval task, mapping blade and vein series data into a shared 128-dimensional latent embedding space. The model’s input utilized an 8-channel tensor (8×512×512), stacking the four ECT maps and four binary masks corresponding to the four nodes of the respective component (blade or vein). The model architecture featured three shared convolutional layers followed by max-pooling and a final linear projection head. Training was conducted using the Contrastive Loss function (margin=1.0), which included a critical step of Hard Negative Mining within each batch to maximize feature discrimination by focusing the loss calculation on the most similar negative pair. The training regime utilized the Adam optimizer (learning rate=10^-4^) over 150 epochs, incorporating a fixed seed (42) data split (80% training, 20% validation) and an Early Stopping mechanism (patience=15, min_delta=0.001) based on the average validation mean Average Precision (mAP). Model performance was evaluated on the validation set using standard retrieval metrics, including mean Average Precision (mAP) and Recall@K (K=1, 5, 10), calculated across both blade-to-vein and vein-to-blade retrieval directions. Following training, the model with the best mAP was used to generate a complete set of 128-dimensional embeddings for all data groups. The structure of the learned latent space was visually inspected using t-SNE.

## 3 Results

### 3.1 The blade and vasculature each contain sufficient information to predict the other

Semantic segmentation models are a Convolutional Neural Network (CNN) in which identity is assigned to each pixel in an image. In our previous work, we generated a high-resolution morphospace of realistic grapevine leaves, where each leaf is composed of two contours representing the blade and vasculature (21). Given the availability of this data, we asked if it was possible to predict the blade from the vasculature, or vice versa. An obvious problem is that both the blade and vasculature are represented as outlines, or with respect to images, as shape masks. The contours of each fall outside the bounds of the other. The Euler Characteristic Transform (ECT) is backed by mathematical proof to distinguish any data object from another, that there is a one-to-one correspondence between ECT and the original data (28). Beyond the ability to extract information from the shape of data objects, the ECT has desirable attributes of featurizing contours represented as embedded graphs in a way that is compatible with CNNs (25). By transforming the ECT into polar coordinates, the ECT features spatially correspond with those of the original shape mask of the contour and are more fully represented throughout the image and accessible to the CNN.

We trained semantic segmentation models, either using the radial ECT and shape mask of the blade to predict vein mask pixels or, reciprocally, using the radial ECT and shape mask of the vasculature to predict the blade mask pixels. Visually, both models reasonably predict the target features (**Figure 1**).

**Figure 1:**
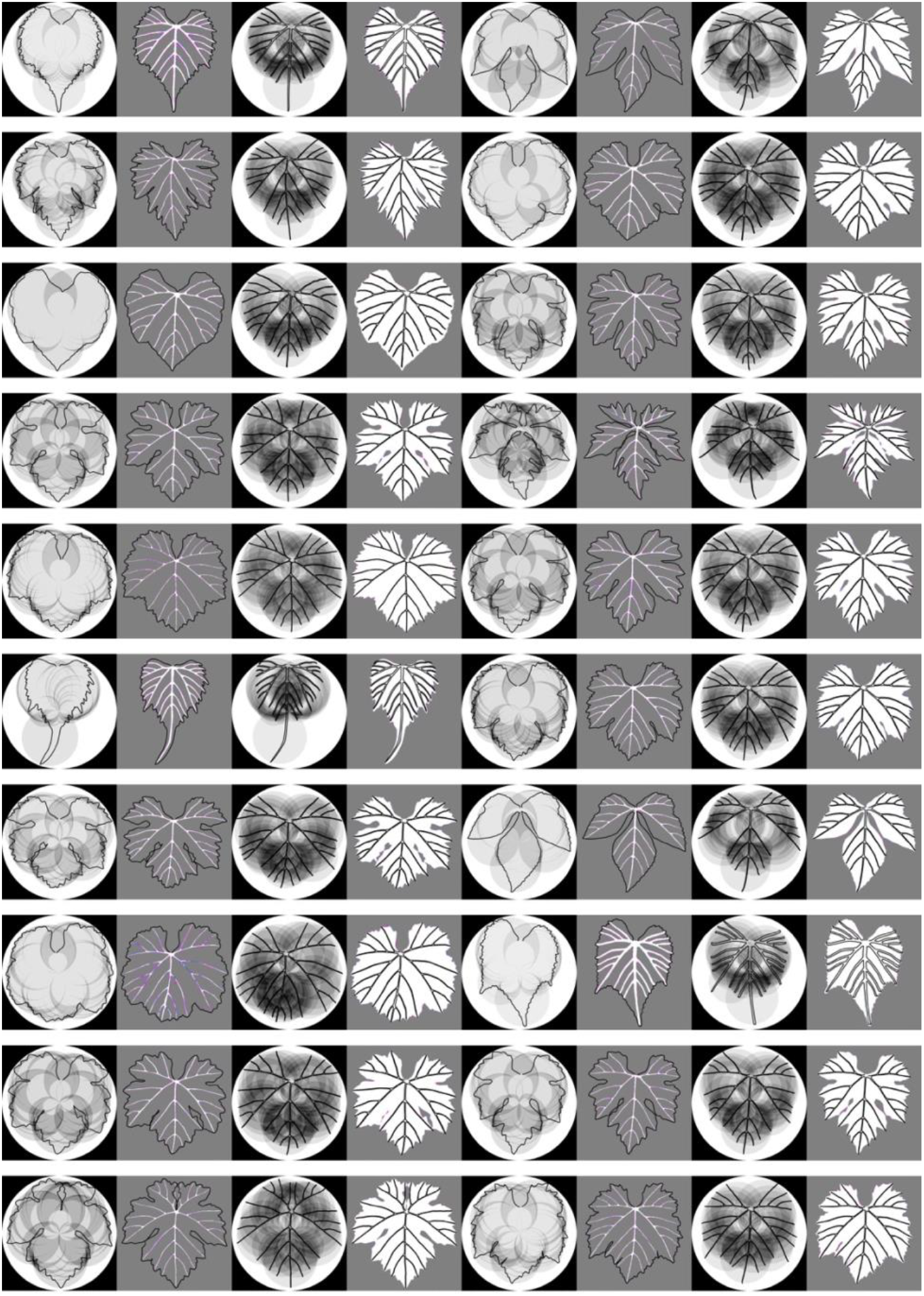
The Euler Characteristic Transform (ECT) and mask of the blade predict the vasculature and vice versa. For each pair of columns, the radial ECT and shape mask input is shown to the left (black background) and the predicted output to the right (gray background). In the prediction on the right, the original input contour is shown as an outline and predicted pixels are colored as true positives (white), false positives (magenta), or false negatives (blue). Two leaves are shown per a row, one leaf in the left four columns and another leaf in the right four columns. For each leaf, the two left columns show prediction of the vasculature from blade and the two right columns the prediction of blade from vasculature.

Specifically, the vasculature, consisting of narrow and intricate veins, is usually predicted with the five major veins and their branches intact, but often has a high number of false positives that represent “near misses” of these structures that are difficult to precisely predict. The Dice coefficient (a measure used to predict the similarity between the ground truth and the prediction sets, ranging from 0 for no overlap to 1 for perfect overlap) reflects this, with the prediction of blade from vasculature achieving a maximum Dice validation score of 0.99 and the prediction of veins from blade achieving a maximum Dice validation score of 0.79. Importantly, this asymmetry in Dice scores reflects differences in geometric complexity and pixel sensitivity between blade and vasculature representations, rather than evidence for a directional or causal relationship between venation and blade patterning. Evaluating the visual results (**Figure 1**) together with Dice coefficient values, and with knowledge of the difficulty of predicting the vasculature to exact pixels, we conclude that both the blade and vasculature of a leaf contain sufficient information to predict the other.

### 3.2 Discriminating one-to-one and many-to-one blade-vasculature relationships

Given that embedded in either the contours of the blade or vasculature is sufficient information to reproduce the other, we hypothesized that there is a one-to-one mapping of blade to vasculature and vice versa. That is, each can predict the other because for a given instance, there is only one possible corresponding match. Assuming that a one-to-one mapping between blade and vein exists, we sought to create a set of leaves with many-to-one relationships. Thin plate splines (TPS) is an interpolation method that resists bending with a penalty, creating a smooth warp between two sets (22). In our data, terminal points of the vasculature correspond with points in the blade, allowing us to warp a blade contour to that of the vein (**Figure 2A**) or vice versa (**Figure 2B**) using these coincident points by TPS. Compared to real leaves (which we call the “matched” dataset) that we assume have a one-to-one correspondence between blade and vein, this easily allows us to create artificial many-to-one datasets for comparison (**Figure 3**).

**Figure 2:**
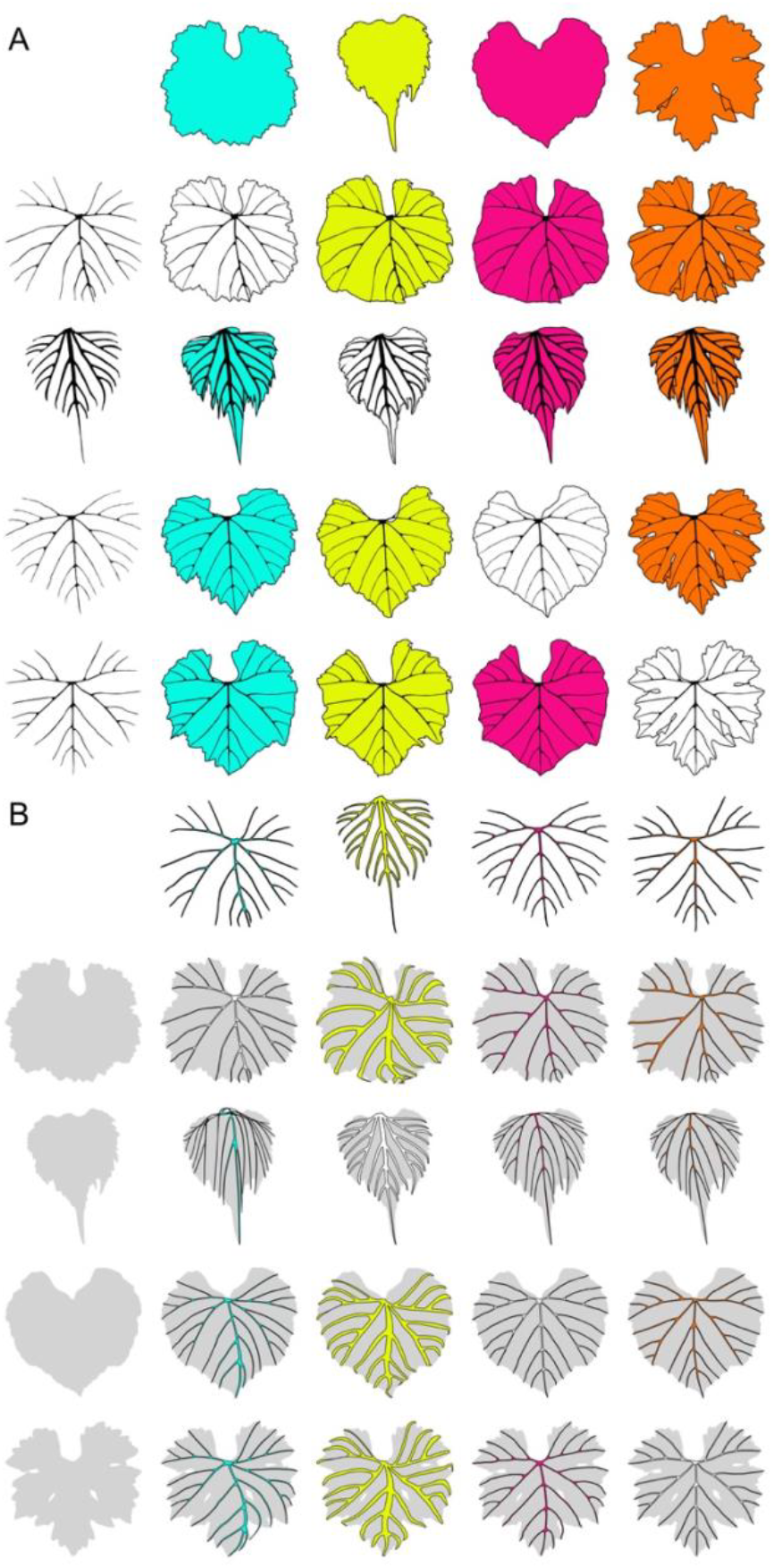
Thin Plate Splines (TPS) swaps blade and vein shapes in a geometrically constrained manner. **A)** Blades (columns, each a different color) warp to constant veins (rows, in black) and **B)** veins (columns, each a different color) warp to constant blades (rows, in gray). For each, the diagonal column represents the original combination of vein and blade for each leaf. All combinations of blade warped to vein or vein warped to blade are shown.

**Figure 3:**
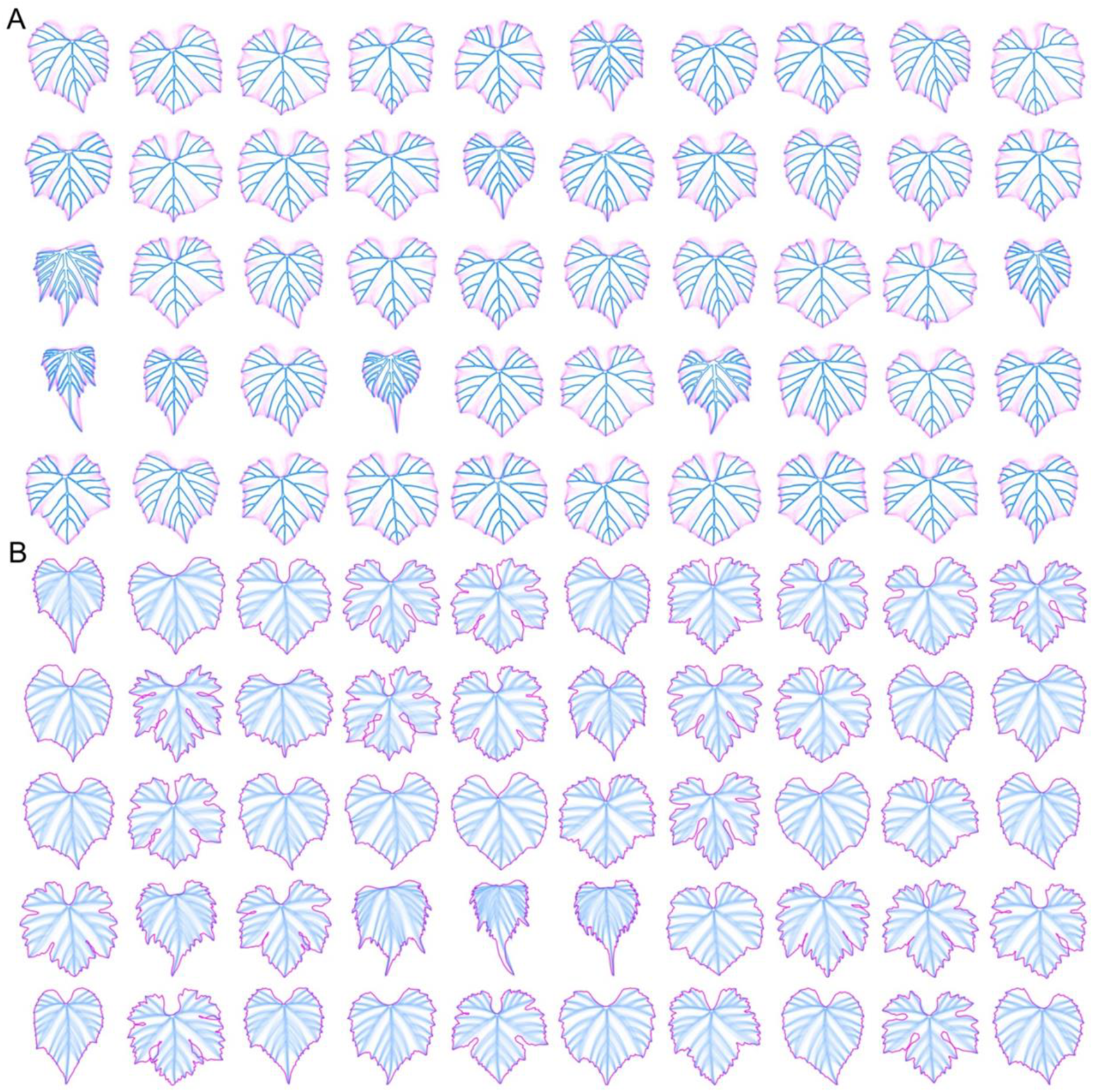
Experimental design to artificially create many-to-one relationships between blade and vein. **A)** For each of 50 anchor veins (blue), the 50 other blades (magenta) are warped. **B)** For each of 50 anchor blades (magenta), the 50 other veins (blue) are warped. In each case for 50 instances of the base component, the 50 other components are warped generating 2,500 synthetic leaves that have a many-to-one relationship.

A two tower CNN is a powerful framework that combines two different data sources, or modalities, and learns to minimize the distance between true pairs (and maximize the distance between non-matches) to place the two different modalities in a common embedding space (29). Blade and vein ECT and shape masks can be thought of as two separate modalities that are paired. We trained a two tower CNN to learn the pairing between blade and vein samples in common embedding space (**Figure 4**). Looking at measures of global differences, there appears to be no differences between the “matched”, “warped blade”, and “warped vein” datasets, suggesting that the TPS warping creates geometrically plausible leaves. This is reflected in the Euclidean distance between pairs in the common embedding space (**Figure 4A**) and that the distribution of the datasets in PCA morphospace largely overlap (**Figure 4B**). The global similarity of warped leaves persists across PCs, but a Linear Discriminant Analysis (LDA) model of dataset as a function of all PCs clearly separates the groups (**Figure 5**), indicating that underlying global similarities are local differences.

**Figure 4:**
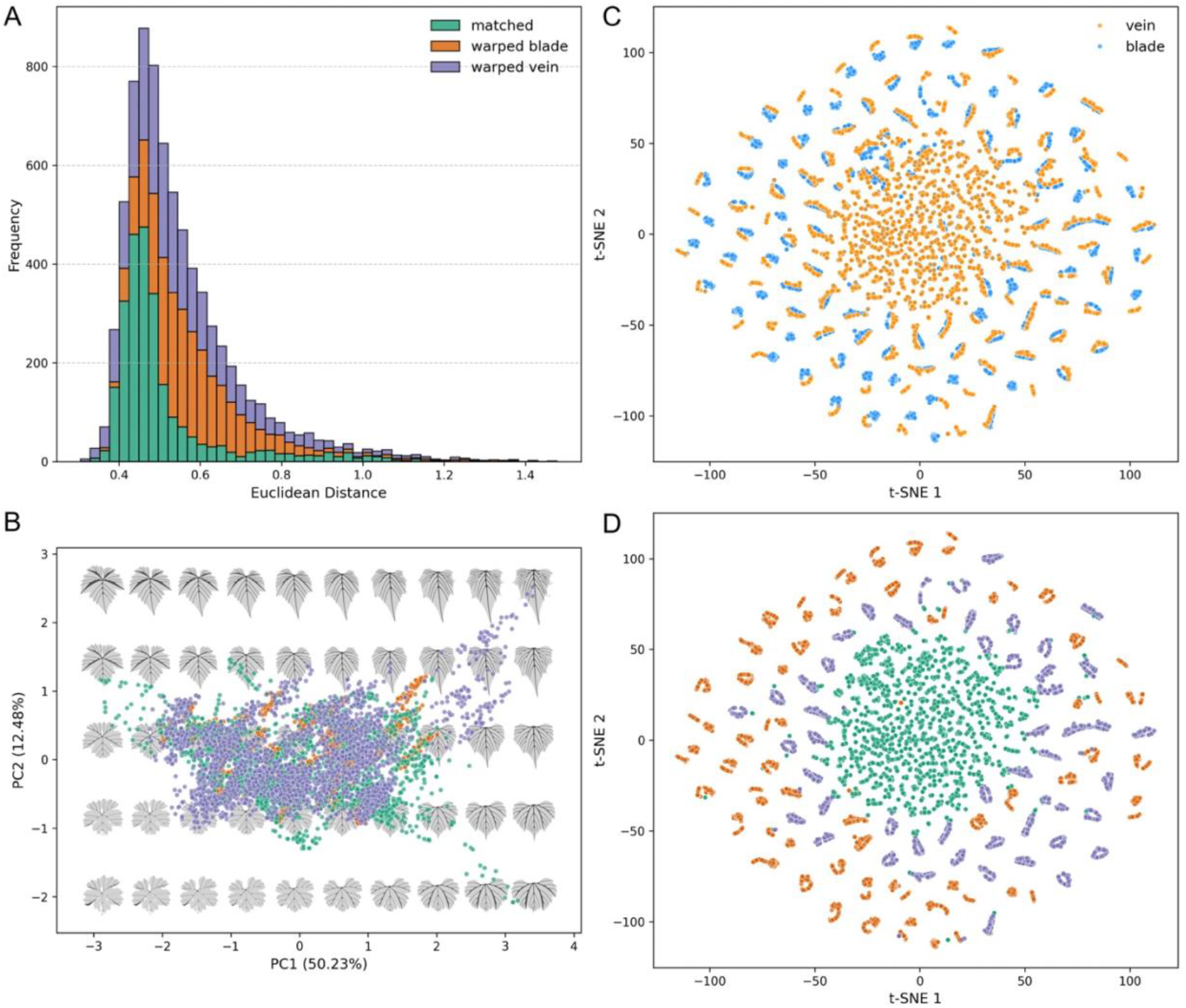
In a common embedding space between blade and vein, swapped leaves are geometrically indistinguishable from true matches at a global level but are different locally. **A)** Histogram of Euclidean distances between blade and vein pairs. Distances in matched, warped blade, and warped vein datasets largely overlap. **B)** A Principal Component Analysis (PCA) of leaf shapes (with eigen leaf shapes in the background) shows large overlap between datasets. **C)** t-SNE on vein and blade embeddings. Vein-blade pairs in the center are so close that there is overplotting, while pairs in the periphery are more separated. **D)** The same t-SNE but colored by dataset, showing the original matched dataset in the center and warped blade and warped vein datasets in the periphery.

**Figure 5:**
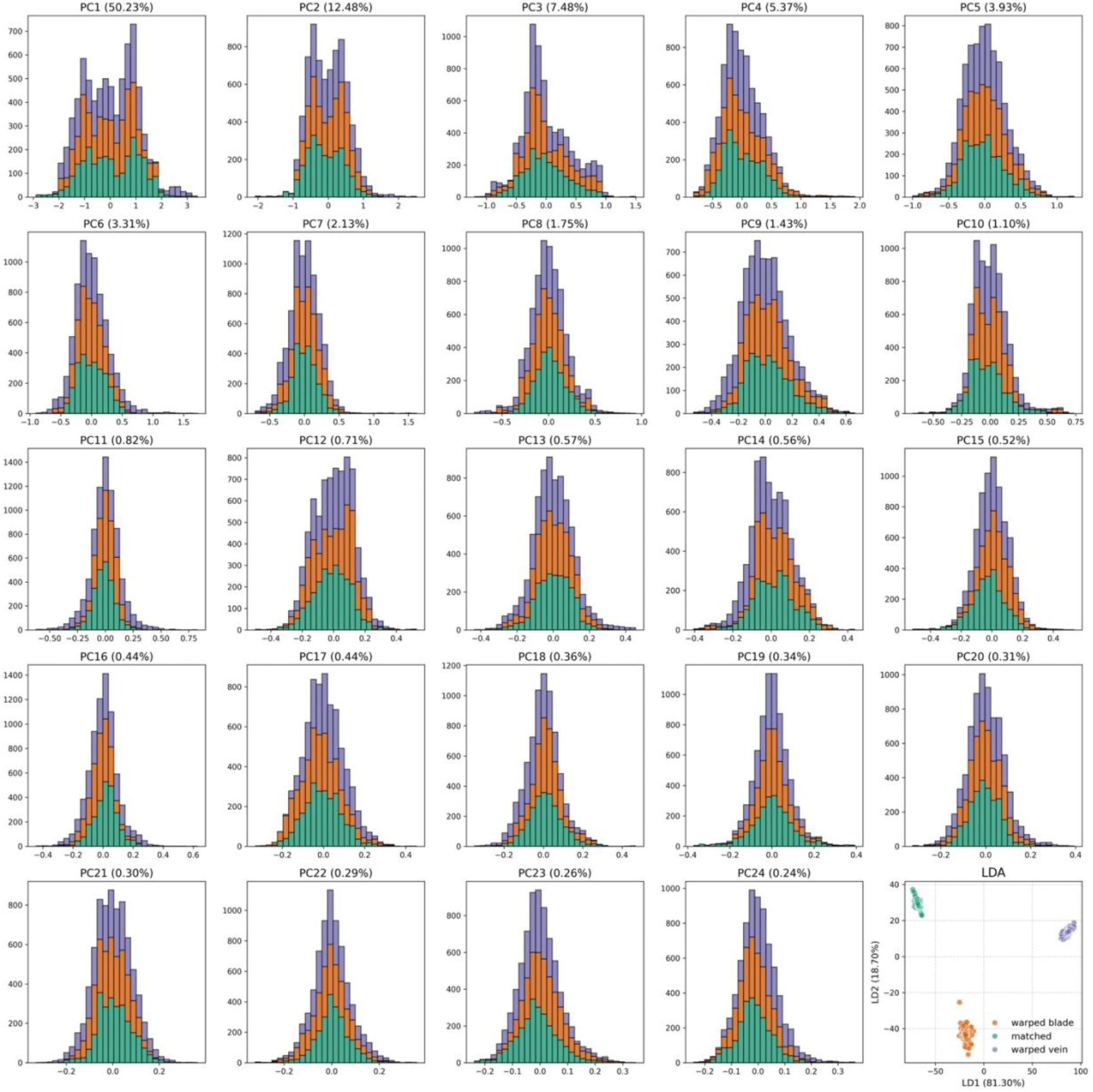
Principal Component Analysis (PCA) fails to distinguish, and Linear Discriminant Analysis (LDA) distinguishes, matched and mismatched leaves. A histogram for each of the first 24 PCs demonstrates that the classes of matched, warped blade, and warped vein largely overlap with each other. In the bottom righthand corner, a LDA performed on all PCs clearly separates the classes.

Despite the geometric plausibility of the TPS warped leaves, non-linear dimension reduction of the resulting two tower CNN embeddings shows clear differences between one-to-one “matched” and many-to-one “warped blade” and “warped vein” datasets. The ability of the two tower CNN to match blade and vein in “matched” data is so strong that they appear overplotted, whereas for the “warped blade” and “warped vein” data there is visible separation between blade and vein (**Figure 4C**), and the “matched” data concentrates in the center of the space whereas the many-to-one datasets at the periphery (**Figure 4D**). We conclude that TPS warping of leaves to create many-to-one datasets are indistinguishable from real leaves with a one-to-one relationship at a global level, but that indeed differences exist at a local level when placed into a common embedding space.

### 3.3 The coupling between blade and vein is determined by the developmental series, not species differences

Leaf shape manifests in a developmental and evolutionary context, both ontogenetically and heteroblastically, and across species. We created intraspecies and interspecies swaps between blade and vein using leaf data from the first four nodes of ten *Vitis* spp. (**Figure 6A**). In intraspecies swaps, the order of the vein and blade across the first four nodes is reversed relative to the other, and either the blade warped to vein (**Figure 6B**) or the vein warped to blade (**Figure 6C**) using TPS. For each of the “warped blade” and “warped vein” series, we take the developmental series in each order. We term “with the series” the order that goes in the correct direction of the unwarped component, and “against the series” the order that goes against the true direction of the unwarped component. Similarly, we swapped vein and blade series between all species combinations, and either warped the blade to vein (**Figure 7A**) or the vein to blade (**Figure 7B**). Unlike the intraspecies species swaps where order could be in either direction, a single order is maintained, even though there is a warped and an unwarped species in each swap.

**Figure 6:**
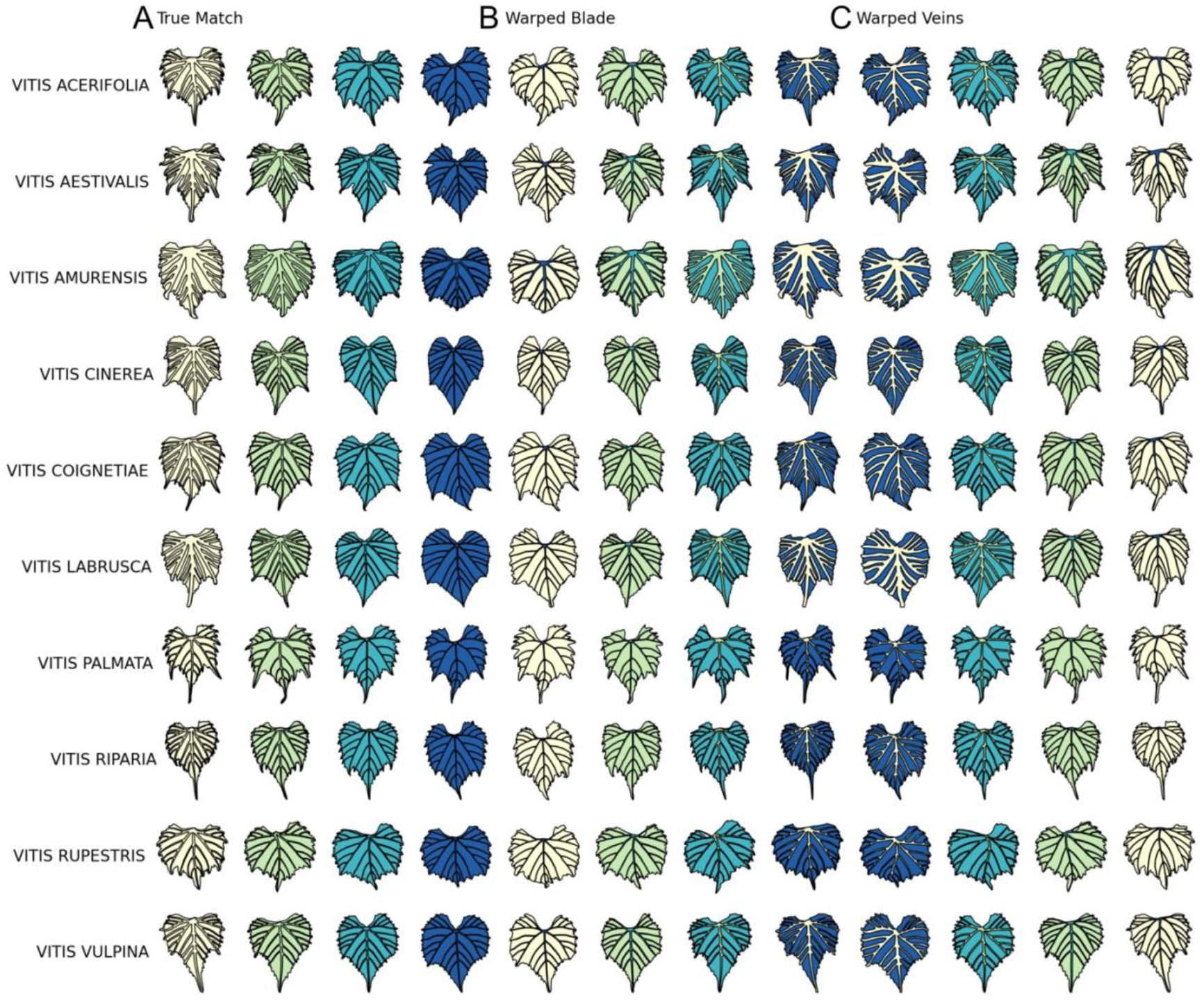
Intraspecies swapping the order of the developmental series between blade and vein. **A)** For each of ten species, the first four, scaled leaves from the tip of the shoot (left to right). Leaves are colored from white to dark blue across the series. **B)** The order of the veins relative to the blade has been swapped and the blade warped to vein. Note the leftmost leaf has the blade of the first node (white) warped to the veins of the fourth node (dark blue). **C)** Again, the order of the veins relative to the blade has been swapped, but the vein warped to blade. For each of the sets of warped blade and warped veins, each possible order (left to right or right to left) is taken. When the order of the unwarped component is in the correct order, it is “with the series”, and when in the reserve order, “against the series”. In each case of the warped datasets, “with the series” is right-to-left and “against the series” is left-to-right.

**Figure 7:**
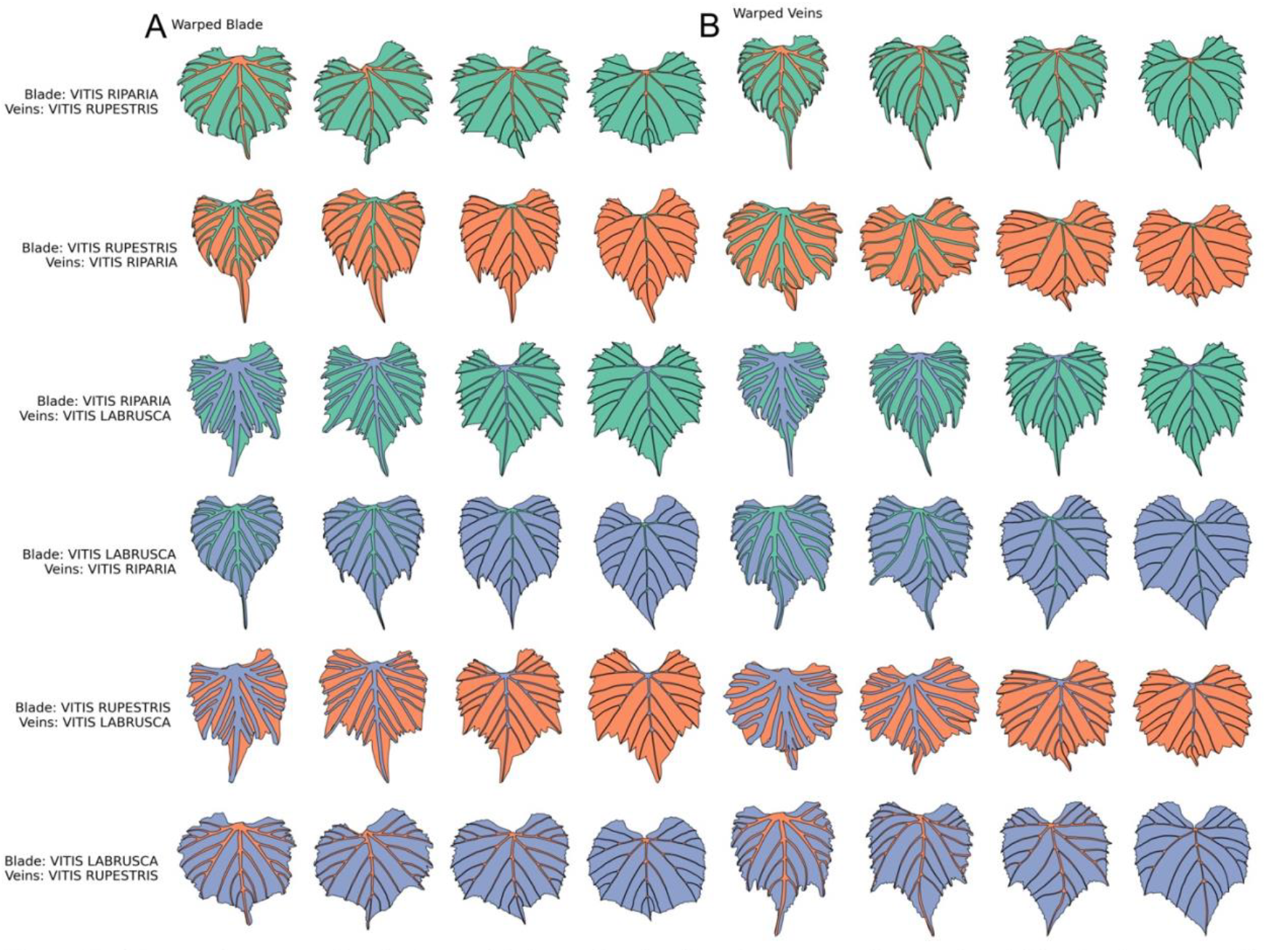
Interspecies swapping between blade and vein. For three example species (*Vitis riparia*; green, *V. rupestris*; orange, and *V. labrusca*; lavender) examples are shown for each combination of blade and vein between species where **A)** the blade is warped to vein and **B)** the vein is warped to blade.

Performing a two tower CNN to create a common embedding space of vein and blade developmental series, we see more obvious separation between vein and blade pairs arising, again, from the many-to-one relationships we have created by swapping components of real leaves (**Figure 8A**). The vein and blade pairs partition into discrete groups composed of true, matched leaves, intraspecies swaps that warp “with the series”, and interspecies swaps, regardless of the identity of the warped or unwarped species (**Figure 8B**). The one unique group that clusters together away from these other groups is the intraspecies swap that warps “against the series”, demonstrating that the natural developmental series is a fundamental component of the relationship between blade and vein. The remaining clusters are defined by species identity (**Figure 8C-D**). Notably, when the blade and vein pair is part of an interspecies swap, it is the unwarped species, not the warped species, that defines the membership in the cluster. We conclude that the relationship between blade and vein is 1) dictated by the progression of the developmental series that is conserved across species and that 2) this relationship occurs in the context of separate, orthogonal species variation, similar to what we described before in grapevine (18).

**Figure 8:**
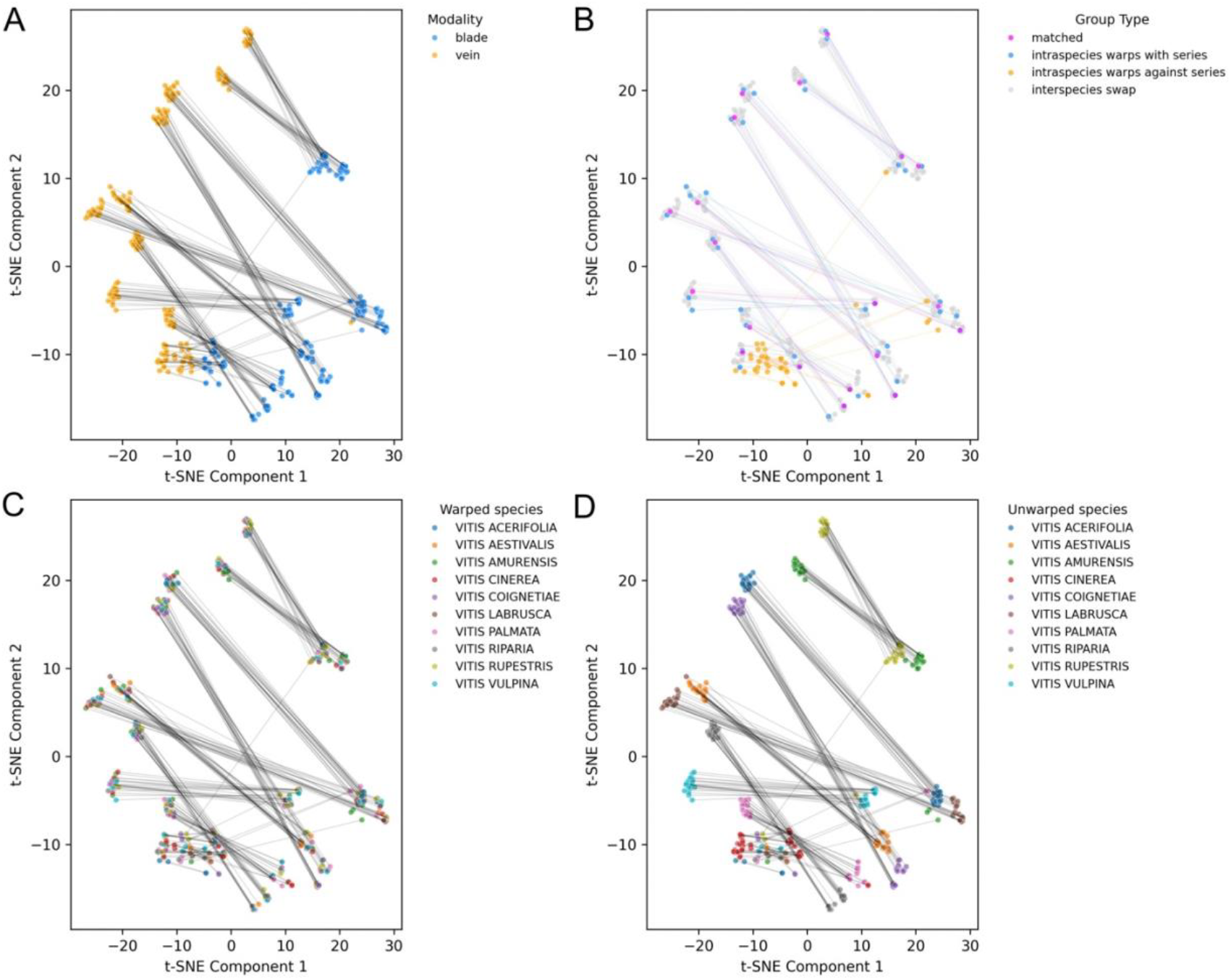
Coupling between blade and vein is determined by the developmental series, not species differences. A common embedding space pairing vein and blade developmental series of the first four nodes (connected by line segments) colored by **A)** vein or blade series, **B)** matched, intraspecies “warps with series” swap, intraspecies “warps against series” swap, and interspecies swap, **C)** warped species interspecies swap, and **D)** unwarped species interspecies swap.

## 4 Discussion

Enabled by the radial ECT that featurizes closed contours into images that spatially correspond with original shape features (25), a semantic segmentation model demonstrates that both the blade and vasculature contain sufficient information to predict the other (**Figure 1**). We take this as an axiomatic observation and the remainder of the analyses in this manuscript are designed to ask why this is the case. We assumed that if there were a many-to-one relationship between blade and veins in leaves that such prediction would not be possible. Assuming that real leaves do have a 1:1 relationship, we purposefully created artificial leaves with many-to-one relationships using TPS (**Figures 2-3**). Remarkably, the set of many-to-one leaves cannot be differentiated from true leaves using global measures of difference, like Euclidean distance or a PCA of leaf shape; yet, using a two tower CNN or an LDA on all PCs, the differences between these sets are obvious (**Figures 4-5**). We interpret this to mean that because our TPS swaps between blade and vein are constrained by coincident points between the vasculature and blade, that individually warped leaves are geometrically plausible, but that such leaves, at an individual level (**Figure 5**) and as a set (**Figure 4D**) are discernable from true leaves by creating mismatched blade and vein pairs and many-to-one relationships, respectively. We also consider the developmental series of the first four nodes of ten *Vitis* spp. and perform intraspecies (**Figure 6**) and interspecies (**Figure 7**) swaps, warping each component to the other and considering all possible orderings. Only reversing the developmental order “against the series” of the unwarped component disrupts the relationship between vein and blade, and the differences in shape between species (defined by the unwarped species) manifest orthogonally and separate from the pairing of vein and blade (**Figure 8**).

Our results help clarify often unspoken assumptions about the relationship between blade and vein. Principal among these is how blade dissection manifests in relation to the vasculature. In *A Practical Ampelography* (10) the vasculature and blade are considered separately. Whereas reniform, orbicular, cordiform, cuneiform, or truncate leaf types are defined by the length and relative angles between the veins, the depth of lobing is considered separately from this, as if vasculature is a chassis on which rests the blade with varying amounts of lobing and dissection. The debate of how to conceptualize the development and evolution of compound leaves mirrors this ambiguity. On one hand there are examples of related species at a microevolutionary level that indeed exhibit varying degrees of dissection, with a continuous path between entire to highly dissected leaves; but on the other hand compound leaves can be viewed as arising through the lens of reactivating indeterminacy and patterning leaflets along primary and secondary axes of the rachis (30). At least in grapevine, our results suggest that there is not a many-to-one relationship; that is, for a given form of the vasculature, there are not varying blade shapes, as depicted in **Figure 2A**, for example. There is empirical evidence for this as well, as we previously showed that losses in blade area resulting from dissection in grapevine are compensated with by longer veins (31). The above makes intuitive sense: if the vasculature and blade are patterned through shared pathways of adaxial-abaxial patterning and auxin (among others), then we expect a constrained relationship in which altering one tissue affects the other. In a mathematical sense, the constraint creates a one-to-one relationship that can be learned by deep learning models.

Beyond the machine-learnable relationships between blade and vein that reveal a one-to-one mapping between their structures (**Figures 4**, **8**), the separate and orthogonal separation by species (**Figure 8**) is intriguing. We previously demonstrated and proposed such a relationship, that despite disparate shape differences between grapevine species, that node-to-node differences in leaf shape arising from development are conserved (18). It will be interesting to apply deep learning-based models predicting vasculature from blade shape (and vice versa) in other taxa and more broadly than we have here. While the relationship between blade and vein is conserved within *Vitis* and *Ampelopsis* spp., we expect at some point for the relationship to fundamentally change, as reflected in qualitatively different vascular patterns seen in leaves with similar shapes across plants. Just as the homology between leaflets and leaf (30), and the possible homology or analogy of the contribution of molecular mechanisms to plant development remain open questions, the extent to which the relationship between blade and vasculature is conserved or diversified across the spectacular variety of leaf shapes remains to be seen.

## Data Availability Statement

Data and code to reproduce this work can be found for the following analyses: Reciprocal prediction of vein and blade from the other, https://zenodo.org/records/16920155; Two tower CNN 1:1 vein:blade relationship, https://zenodo.org/records/17014105; Interspecies and Intraspecies developmental series swaps, https://zenodo.org/records/17013783

## Conflict of Interest Statement

This study was conducted in collaboration with E & J Gallo Winery, which provided access to commercial vineyards throughout California and from which data were collected. Peter Cousins, an author of this study, is an employee of E & J Gallo Winery.

## Acknowledgements

This research was funded by a PRFU project (Code D01N01UN070120220001) approved in 2022 by the Algerian Ministry of Higher Education and Scientific Research, supporting Z. Laiadi and his research team. DHC, EM, and YA were supported by the National Science Foundation Plant Genome Research Program award numbers IOS-2310355, IOS-2310356, and IOS-2310357. ZM was supported by the Canada Research Chairs Program. JFS was supported by an NSF Postdoctoral Research Fellowship under Award No. 2305703.

## Notes

https://zenodo.org/records/16920155

https://zenodo.org/records/17014105

https://zenodo.org/records/17013783

